# Diverse mechanisms of DDX3Y suppression by DDX3X

**DOI:** 10.1101/2025.02.08.637260

**Authors:** Xiaolu Xu, Shuo Wei

## Abstract

The DEAD-box RNA helicase DDX3X has important roles in development and disease. Loss of DDX3X during developmental and pathological processes such as tumorigenesis can lead to compensatory upregulation of the close paralog DDX3Y in males, which may underlie the sexual dimorphism displayed by some DDX3X-associated diseases. However, how DDX3X cross-regulates DDX3Y remains largely unknown. Here, we investigated the regulation of DDX3Y by DDX3X in two male-derived human cancer cell lines, HCT116 and U87MG. Depletion of DDX3X in HCT116 cells results in moderately increased DDX3Y mRNA and protein, in part due to stabilization of *DDX3Y* transcripts. Conversely, reduction of DDX3X in U87MG cells markedly upregulates DDX3Y protein without affecting its mRNA, mainly by enhancing DDX3Y protein stability. We further show that DDX3X physically interacts with DDX3Y. DDX3Y is much less stable than DDX3X in U87MG cells, and substitution of two lysine residues in DDX3Y with the corresponding arginine in DDX3X stabilizes DDX3Y. Thus, the compensatory upregulation of DDX3Y following DDX3X loss can occur at either transcript or protein level, suggesting complex and cell type-specific cross-regulation between these X- and Y-linked paralogs to keep the total DDX3 dosage in check.

## Introduction

The DEAD-box RNA helicase DDX3 functions in nearly all aspects of RNA metabolism, including transcription, splicing, nuclear export, degradation, translation and stress response (Mo *et al*, 2021; Soto-Rifo & Ohlmann, 2013). The human genome contains two functional *DDX3* homologs, the X-linked *DDX3X* and the Y-linked *DDX3Y*, which are believed to have evolved from a single autosomal gene (Bellott *et al*, 2014; Wilson & Makova, 2009). DDX3X is ubiquitously transcribed and translated, and has been associated with various diseases such as birth defects, viral infection, inflammation, and cancer (Gadek *et al*, 2023; Mo *et al*., 2021). In particular, mutations in *DDX3X* cause DDX3X syndrome, a neurodevelopmental disorder with additional defects in the neural crest (Levy *et al*, 2023; Perfetto *et al*, 2021; von Mueffling *et al*, 2024). DDX3X also acts both as an oncogene and a tumor suppressor, and high DDX3X expression can serve as a positive or negative prognostic factor, depending on the cancer type (Lin, 2019). In contrast, while DDX3Y is also actively transcribed in most tissues, its protein is often undetectable outside of the male reproductive system, likely due to post-transcriptional regulations such as inhibited translation (Ditton *et al*, 2004). Loss of *DDX3Y* has been linked to Sertoli cell-only syndrome, a severe testicular defect (Foresta *et al*, 2000).

DDX3X and 3Y share ∼91% identities in protein sequence and are functionally interchangeable in protein synthesis, although differences in stress granule formation and translational repression have been reported (Shen *et al*, 2022; Venkataramanan *et al*, 2021). At cellular level, both human DDX3X and 3Y can rescue the growth arrest of a BHK21 hamster cell line harboring a temperature-sensitive *Ddx3x* mutation (Sekiguchi *et al*, 2004). Interestingly, cells are particularly sensitive to total dosage of DDX3, and perturbation of one paralog results in the opposite response of the other. In addition, increasing copy number of Y chromosome correlates with shorter half-life of *DDX3X* mRNA *in vitro*, implying that DDX3Y negatively regulates *DDX3X* transcript stability (Rengarajan *et al*, 2025). Loss of DDX3X has also been shown to cause compensatory upregulation of DDX3Y *in vivo*. For example, conditional knockout of *Ddx3x* in neural progenitor cells leads to profound brain defects in the homozygous female mice, but the hemizygous males are normocephalic with moderate but significant increase in *Ddx3y* mRNA (Hoye *et al*, 2022; Patmore *et al*, 2020). *DDX3X* mutations have been detected in various types of lymphoma including ∼30% cases of Burkitt lymphoma, which has a 3:1 male vs. female incidence ratio. Loss of DDX3X function prevents *MYC* oncogene-driven lymphomagenesis by moderating global protein synthesis and buffering MYC-induced proteotoxic stress, but established male lymphoma cells can overcome this effect by aberrantly upregulating DDX3Y protein levels (Gong *et al*, 2021; Lacroix *et al*, 2022). Surprisingly, this upregulation is not accompanied by alterations in *DDX3Y* mRNA abundance, and there is no evidence suggesting that DDX3X can directly control DDX3Y translation (Gong *et al*., 2021), raising the question how this cross-regulation is achieved. To understand how DDX3X regulates DDX3Y levels, we performed loss-of-function studies for DDX3X in human male-derived HCT116 and U87MG cell lines. Our results indicate that DDX3X can suppress DDX3Y at either mRNA or protein level in a cell type-dependent manner, providing a possible explanation to reconcile the discrepancies in the literature.

## Materials and methods

### Reagents

Full-length cDNA clones for human *DDX3X* (BC011819) and *DDX3Y* (BC034942) were purchased from GE-Dharmacon and subcloned into a pCS2+ expression vector with a C-terminal HA or FLAG tag. Primers used in the subcloning are listed in Supplementary Table S1. Pharmacological inhibition of transcription was performed with Actinomycin D (Sigma-Aldrich A1410). Primary antibodies used in this study include mouse anti-DDX3X (Santa Cruz sc-81247; 1:1,000), rabbit anti-DDX3X (Bethyl Laboratories A300-474A; 1:1,000), rabbit anti-DDX3X (Aviva OAAB01241; 1:1,000), rabbit anti-HA (Cell signaling Technology C29F4; 1:2,000), rabbit anti-GFP (Invitrogen A-11122,1:1,000) and custom-made rabbit DDX3Y antibody from Dr. Nicholas Grigoropoulos (1:2,000) (Gong *et al*., 2021). Secondary antibodies include HRP-conjugated mouse anti-β-actin (Cell Signaling Technology 8H10D10, 1:5,000), HRP-linked horse anti-mouse IgG (Cell Signaling Technology 7076, 1:10,000) and HRP-linked goat anti-rabbit IgG (Cell Signaling Technology 7074, 1:10,000).

### Cell culture and transfection

U87MG cells (ATCC) were cultured in MEM (Corning) supplemented with 10% fetal bovine serum (FBS; Gibco) at 37°C with 5% CO_2_. HCT116 cells (ATCC) were cultured in McCoy’s 5A (ATCC) supplemented with 10% fetal bovine serum (FBS, Gibco) at 37 °C with 5% CO_2_. Cells were transfected with 10nM DDX3X, DDX3Y or control siRNA (Dharmacon J-006874-06-0002, L-011904-01-0005, and D-001810-01-05) at 50% confluency using Lipofectamine RNAiMAX (Invitrogen), or with plasmids at 70% confluency, using Lipofectamine 3000 (Invitrogen). For experiments that used both siRNA and plasmid, siRNA was transfected first for 24h before plasmids being transfected for another 48h.

### CRISPR/Cas9-mediated genome editing

Small guide RNAs (sgRNAs) were designed to target exon 5 or exon 1 of the DDX3X gene using CRISPRscan. Protospacers DDX3X-g1 and DDX3X-g2 were cloned into a pSpCas9(BB)-2A-GFP/PX458 expression vector (Addgene 48138) using a scarless golden gate cloning strategy as described (Ran *et al*, 2013). Primers used in subcloning the protospacers are listed in Supplementary Table S1. Plasmid was transfected in U87MG cells at 500ng/ml for 48 hours followed by Fluorescence-activated Cell Sorting (FACS) to sort GFP-expressing cells and plated them individually into 96 well plates. Clones were cultured in MEM supplemented with 20% FBS and expanded for 2-3 months to collect enough samples for sequential procedures. Genomic DNA was isolated and then PCR amplified using genotyping (GT) primers followed by PCR cloning (NEB E1202S) and sanger sequencing. Total RNA was isolated using the RNeasy Mini Kit (Qiagen) and then reverse transcribed into cDNA using the iScript cDNA Synthesis Kit (Bio-Rad Laboratories) with DNase I (Qiagen) treatment. With the cDNA as the template, PCR amplification was performed, followed again by PCR cloning and sanger sequencing.

### Western blotting and RT-qPCR analyses

Cells were lysed on ice in RIPA lysis buffer (Fisher Scientific 89901) with Halt protease inhibitor cocktail (ThermoFisher 78429) and Halt phosphatase inhibitor cocktail (ThermoFisher 78420) added, and processed for western blotting as described (Li *et al*, 2018). Immunoprecipitation (IP) was carried out with Pierce™ Anti-HA Agarose (ThermoFisher 26181) at 4°C overnight with the bound protein released by boiling in Laemmli SDS sample buffer (ThermoFisher J61337-AD) at 95°C for 10 min. Blots were detected with HRP-conjugated antibodies and Clarity Western ECL substrate (Bio-Rad #1705061) using a Bio-Rad ChemiDoc touch imager. Total RNA was extracted from collected embryos using RNeasy Mini Kit (Qiagen 74104) and then reverse-transcribed into cDNA using iScript cDNA Synthesis Kit (Bio-Rad 170-8891). Using qMax Green qPCR Mix (Accuris ACC-PR2000-L-100), quantitative PCR (qPCR) was carried out on Quant Studio 6 Flex Real-Time PCR system (Applied Biosystems). The cycling conditions include an initial denaturation step at 95°C (2 min), 40 cycles of 5 sec at 95°C and 30 sec at 60°C. Primers used for RT-qPCR are listed in Supplementary Table S1.

## Results

### Loss of DDX3X upregulates DDX3Y mRNA and protein in HCT116 cells

Previous studies have shown the upregulation of *Ddx3y* mRNA upon knockout of *Ddx3x* in mice (Hoye *et al*., 2022; Patmore *et al*., 2020). However, it remained unclear how the translated Ddx3y protein is affected, due to the lack of an antibody that can differentiate between Ddx3x and 3y. With a recently generated antibody that specifically recognizes human DDX3Y but not 3X (Gong *et al*., 2021), we were able to use cultured human cell lines to examine the regulation of DDX3Y protein by DDX3X. In HCT116 colon cancer cells, we detected 44% increase in endogenous DDX3Y following siRNA-mediated knockdown of DDX3X (Figure 1A). This was accompanied by 98% increase in *DDX3Y* mRNA (Figure 1B), suggesting that the upregulation of DDX3Y protein was primarily caused by elevated transcripts. As DDX3X has been shown to function in various processes of RNA metabolism, including transcription, nuclear export and stability (Mo *et al*., 2021; Soto-Rifo & Ohlmann, 2013), we assessed if the stability of *DDX3Y* mRNA can be altered by DDX3X. After treating the cells with the transcription inhibitor actinomycin D, we measured the amount of *DDX3Y* mRNA over time (Figure 1C). Upon knockdown of DDX3X, *DDX3Y* mRNA was degraded more slowly with a half-life of 4.3 hr as compared to control, which had a half-life of 3.0 hr (Figure 1D). In addition, after 8-hr treatment with actinomycin D, there was a significant difference in the fold change of *DDX3Y* mRNA between control and DDX3X knockdown cells (Figure 1E), implying that DDX3X destabilizes *DDX3Y* mRNA. However, we cannot rule out the possibility that DDX3X also represses *DDX3Y* transcription.

**Figure 1.**
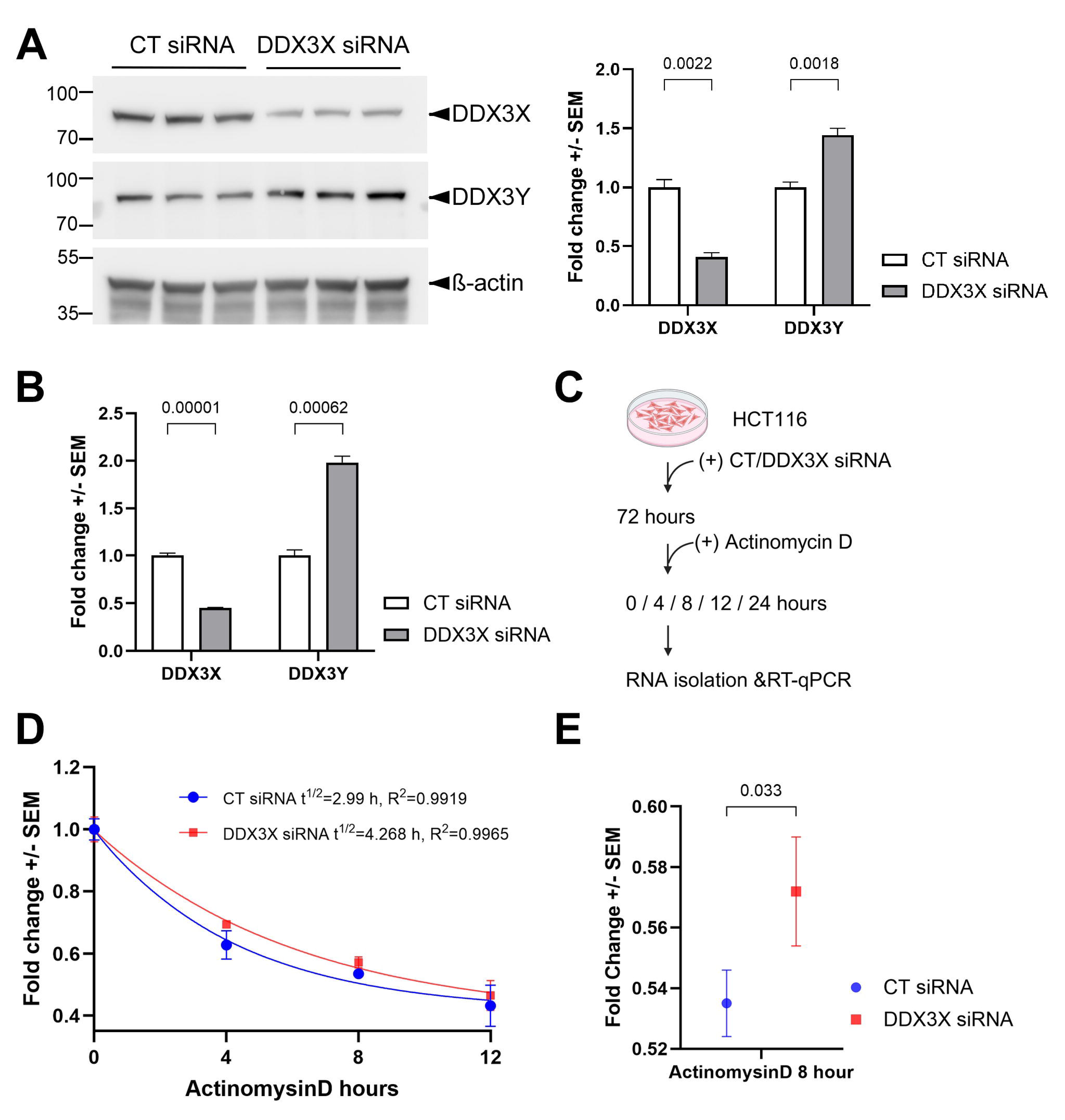
Loss of DDX3X leads to enhanced levels of DDX3Y mRNA and protein in HCT116 cells. **(A)** HCT116 cells were transfected with indicated siRNAs, and cell lysates were processed for western blotting with the indicated antibodies. Representative blots of three biological replicates are shown on the left, and quantification on the right. **(B)** The same batch of cells from **(A)** was processed for RT-qPCR for the indicated mRNAs. **(C)** Flowchart of the mRNA degradation assay. **(D)** Degradation curve of *DDX3Y* mRNA. Fold changes of mRNA at indicated time points post-treatment as compared with 0 hr were calculated and fitted to one phase exponential decay function. **(E)** Fold changes of *DDX3Y* mRNA from control and DDX3X knockdown cells after 8 hr of actinomycin D treatment. All experiments were performed with three biological replicates, and results shown are fold change +/-SEM with p values calculated by unpaired student t-tests.

### CRISPR/Cas9-introduced deletions at the *DDX3X* locus result in reduced but not eliminated DDX3X protein expression in U87MG cells

We next turned to U87MG, a glioma cell line. To knock out *DDX3X* in these cells, we used two guide RNAs (gRNAs) individually for CRISPR/Cas9-mediated genome editing and obtained one clone with each gRNA (Supplementary Figure S1). Sequencing of both the genomic DNA and the cDNA indicated that Clone 1 (obtained with gRNA-1) contained a 159-bp in-frame deletion of the entire Exon 5 in the coding sequence of the mRNA, which resulted in a smaller DDX3X protein with residues 96-148 deleted (Supplementary Figure S2). In contrast, Clone 2 (obtained with gRNA-2) had a 14-bp deletion that was confirmed by sequencing of the *DDX3X* cDNA (Figure 2A). Although our initial western blotting with a Bethyl anti-DDX3X antibody (raised against residues 1-50) suggested that *DDX3X* was completely knocked out in Clone 2, another antibody from Santa Cruz (raised against the broader N-terminus region) picked up a band with a size similar to that of DDX3X but a significantly lower intensity (∼34% of parental cells; Figure 2B). We originally suspected that this band represented DDX3Y, as the two paralogs share high sequence homology. However, this band was reduced when the cells were transfected with an siRNA targeting *DDX3X* but not another one targeting *DDX3Y* (Figure 2C), indicating that it represented DDX3X. A careful re-examination of the *DDX3X* cDNA revealed that the cells likely utilized a slightly downstream and originally out-of-frame AUG to restore the open reading frame after the deletion (Figure 2A), thereby retaining most of DDX3X protein sequence and probably function. This suggests the existence of selection pressure against loss of DDX3X function. Indeed, a search against the DepMap portal (Tsherniak *et al*, 2017) did infer a negative impact of DDX3X loss on U87MG cells (Supplementary Figure S3). The lower endogenous DDX3X protein levels in Clone 2 are probably due to translational repression by the upstream AUG, the original start codon that became out of frame in this clone (Figure 2A). A similar translational repression by upstream AUGs has been reported for DDX3Y (Jaroszynski *et al*, 2011).

**Figure 2.**
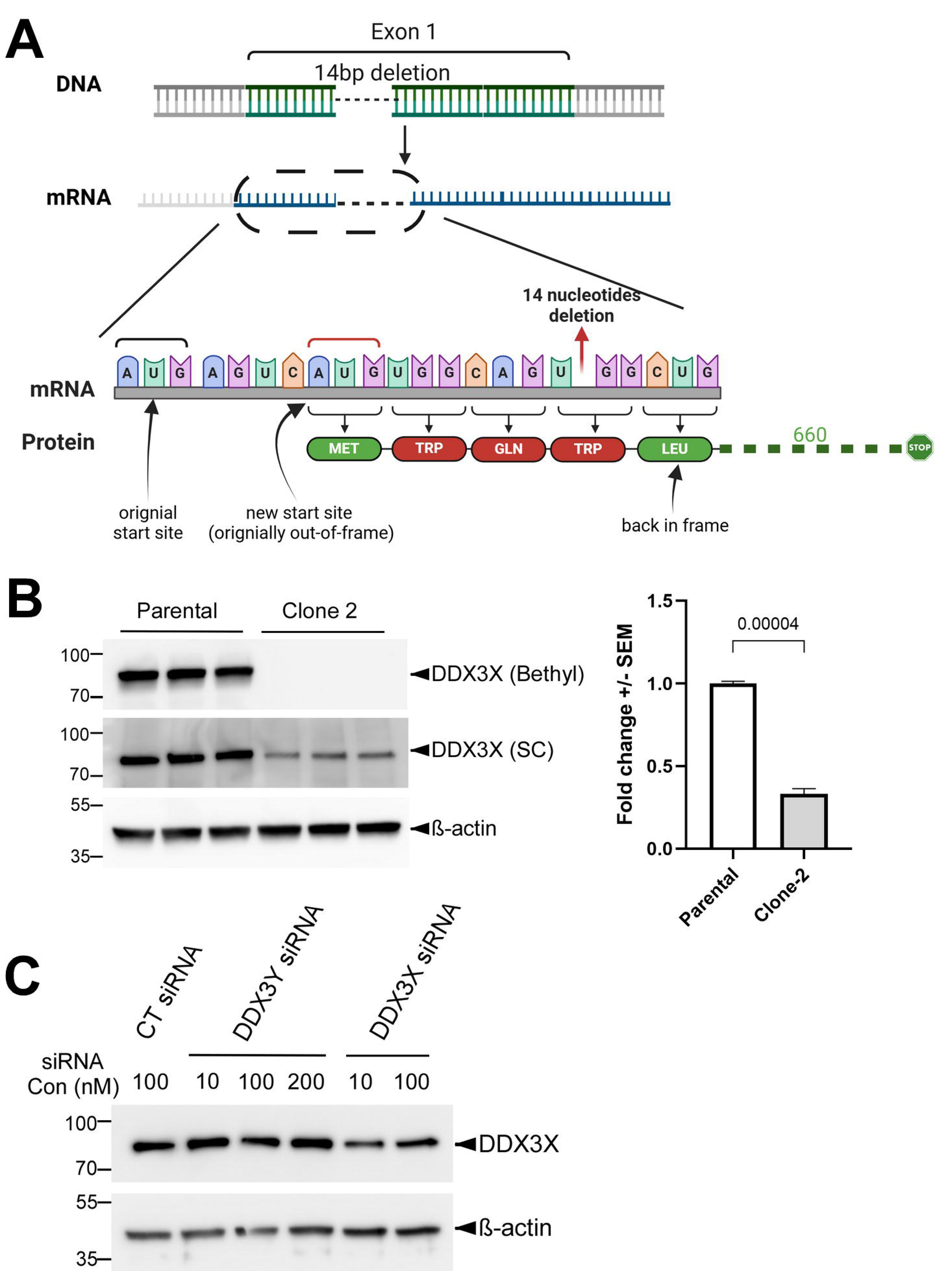
U87MG Clone 2 cells generated by CRISPR/Cas9 express reduced DDX3X protein. **(A)** Diagram of the genotyping results of Clone 2 and the putative encoded DDX3X protein. After the deletion of 14 nucleotides caused by CRISPR/Cas9, which was confirmed at both genomic DNA and cDNA levels, Clone 2 cells likely utilized a downstream translation start site to restore the open reading frame and prevent a complete loss of DDX3X protein. Residues in red were introduced by the mutation, while those in green were unchanged. **(B)** Representative western blots showing endogenous DDX3X in parental U87MG and Clone 2 cells (three biological replicated each), obtained using anti-DDX3X antibodies from two different sources. Quantification of the DDX3X band intensity detected with the Santa Cruz (SC) antibody is shown on the right. **(C)** Clone 2 cells were transfected with the indicated concentration of control (CT), DDX3X and 3Y siRNA, and cell lysates were processed for western blotting with the SC antibody.

### DDX3X promotes DDX3Y protein turnover in U87MG cells

Since the endogenous DDX3Y was not detectable in either Clone 2 or the parental U87MG cells with the specific antibody used in Figure 1A, we transfected these cells with a plasmid encoding C-terminally HA-tagged DDX3Y. Western blotting with an anti-HA antibody showed higher levels of the exogenous DDX3Y protein with a fold change of ∼6.9 in Clone 2, as compared to parental cells transfected with the same amount of plasmid (Figure 3A). As the RNA-binding and helicase domains of DDX3X remain intact in Clone 2, it is likely that this effect was caused by reduced DDX3X levels in this clone. To confirm this, we knocked down DDX3X in the parental U87MG cells and also observed a significant increase in the exogenous DDX3Y protein as compared to control (Figure 3B). RT-qPCR result of mRNA extracted from the same batch of cells indicated that cells expressing exogenous DDX3Y contained 300-fold more *DDX3Y* mRNA than mock-transfected cells (data not shown). Thus, the total *DDX3Y* mRNA essentially represented the exogenous transcripts. No significant change in *DDX3Y* mRNA was found upon DDX3X knockdown (Figure 3C), suggesting that DDX3X regulates DDX3Y at protein level, such as translation or turnover. To examine the potential effect of DDX3X on DDX3Y turnover, we treated cells with the translation blocker cycloheximide (CHX) and monitored the degradation of exogenous DDX3Y. In parental U87MG, DDX3Y protein became undetectable within 4 hr of CHX treatment. By contrast, in Clone 2 cells, where there was reduced endogenous DDX3X, degradation of DDX3Y protein was greatly slowed down (Figure 3D, E). Interestingly, DDX3X was much more stable despite the high sequence homology to DDX3Y, with over 80% of protein remained even after 12 hr of CHX treatment (Figure 3D, E). These results indicate that DDX3X facilitates the turnover of DDX3Y protein in U87MG cells.

**Figure 3.**
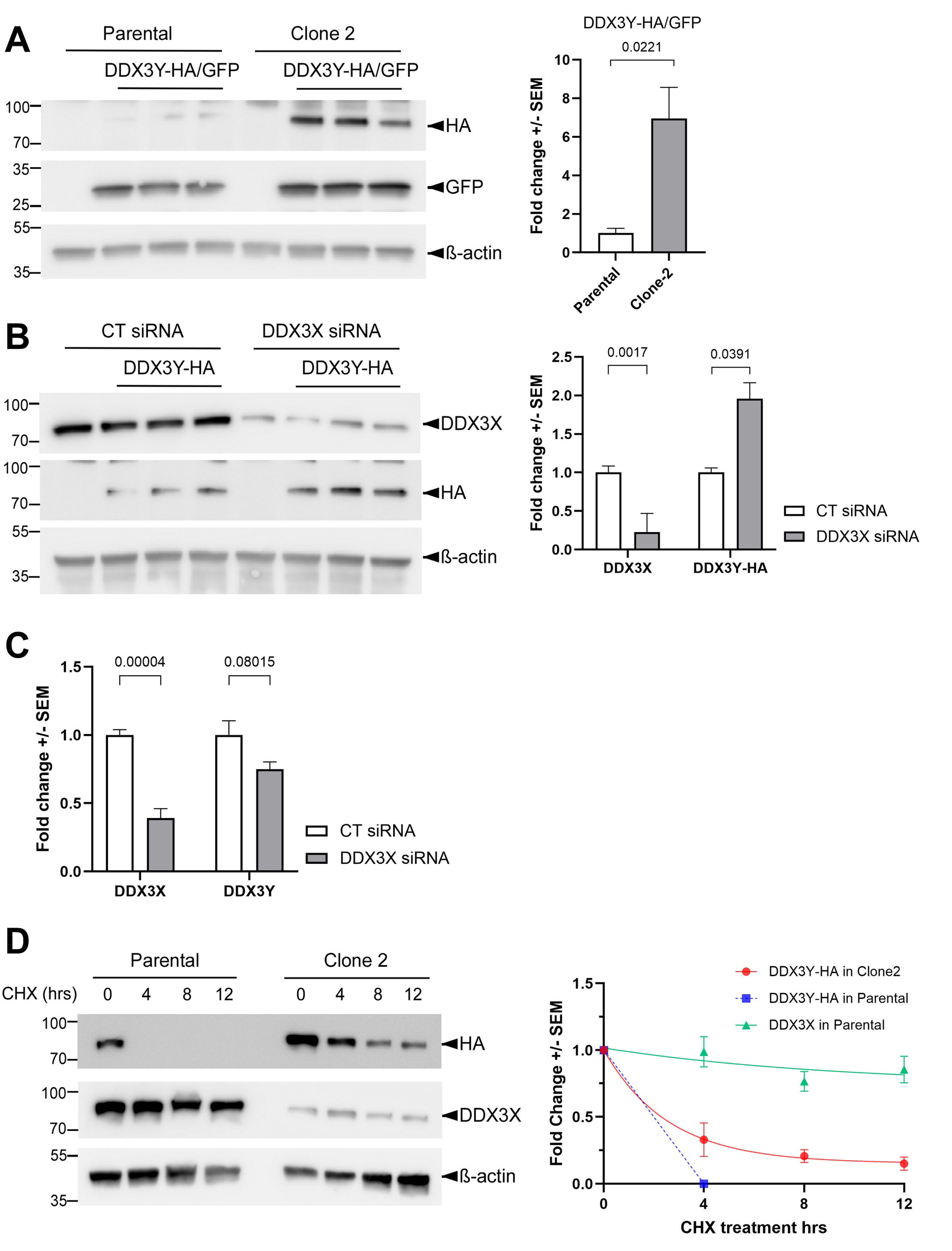
DDX3X facilitates DDX3Y degradation in U87MG cells. **(A)** Parental U87MG and Clone 2 cells were transfected with empty vector or a plasmid encoding HA-tagged DDX3Y; a second plasmid encoding GFP was co-transfected as a control for transfection efficiency. Western blotting for cell lysates was performed with the indicated antibodies, and DDX3Y-HA levels normalized against GFP are shown on the right. **(B)** Parental U87MG cells were transfected with the indicated siRNAs and a plasmid encoding DDX3Y-HA. Western blots for cell lysates using the indicated antibodies are shown on the left, and quantification of endogenous DDX3X and exogenous DDX3Y-HA on the right. **(C)** RT-qPCR results of the indicated mRNAs in the same batch of cells from **(B)**. **(D)** Parental U87MG and Clone 2 cells were transfected to expressed DDX3Y-HA before being treated with 20 μM cycloheximide. Cells were collected at the indicated time points post-treatment and processed for western blotting. Degradation curves of endogenous DDX3X and exogenous DDX3Y-HA on the right were plotted using fold changes of proteins as compared with 0 hr, followed by fitting to one phase exponential decay function. All experiments were performed in three biological replicates, and results shown are fold change +/-SEM with p values calculated by unpaired student t-tests.

### DDX3X interacts with DDX3Y, which contains two lysine residues contributing to the lower stability

Both DDX3X and the yeast homolog Ded1p have been shown to form oligomers to unwind RNA duplexes cooperatively (Putnam *et al*, 2015; Song & Ji, 2019). Based on the high sequence homology between DDX3X and 3Y, we hypothesized that these two paralogs interact with each other. To test this hypothesis, we co-expressed C-terminally HA-tagged DDX3Y and FLAG-tagged DDX3X in Clone 2 cells and performed co-IP using agarose beads conjugated with an anti-HA antibody. As shown in Figure 4A, Flag-tagged DDX3X was pulled down along with the HA-tagged DDX3Y. Likewise, endogenous DDX3X also co-precipitated with HA-tagged DDX3Y in parental U87MG cells (Figure 4B), further confirming the binding of DDX3X to DDX3Y.

**Figure 4.**
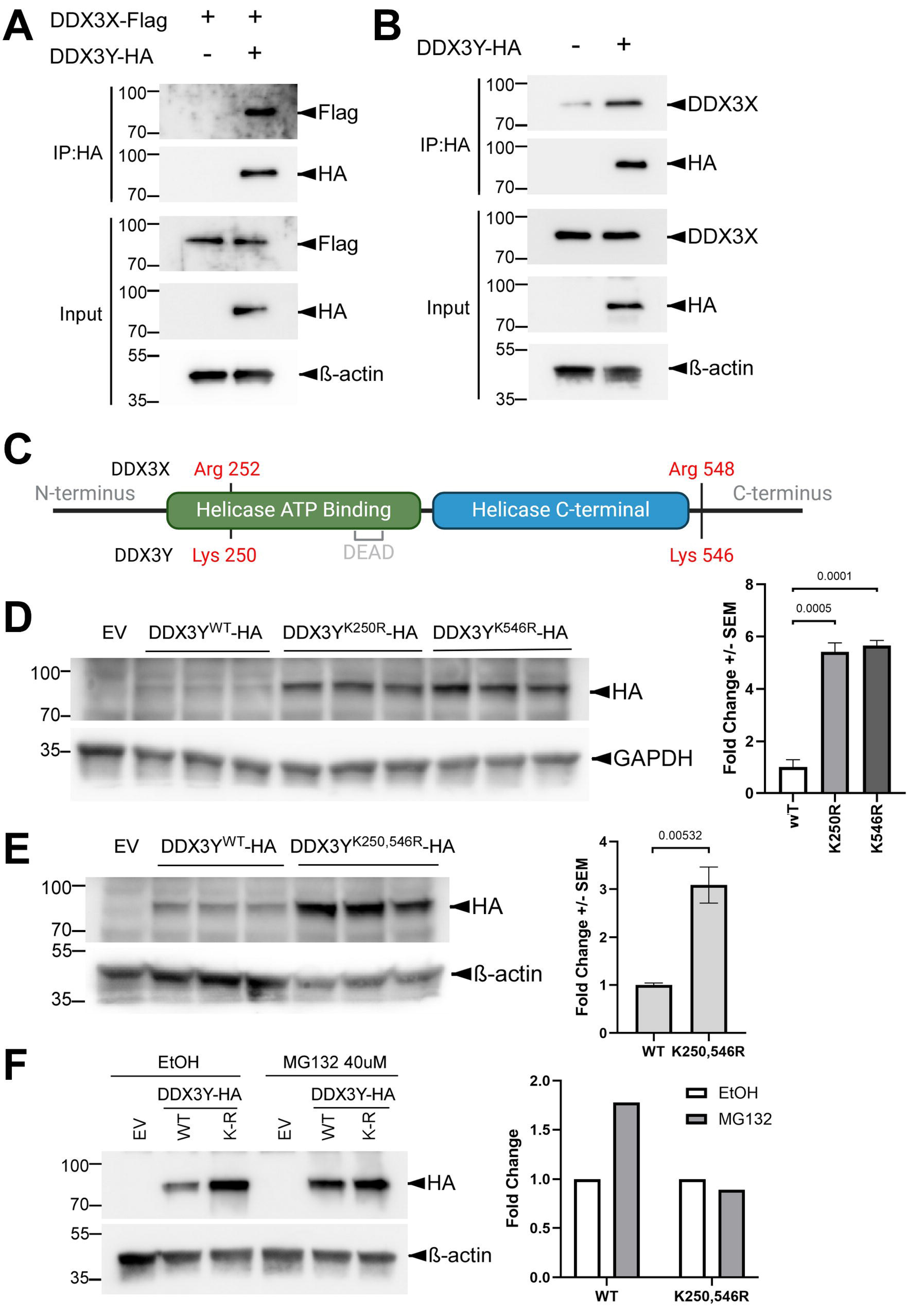
DDX3X binds DDX3Y, and substituting the lysine residues by arginine stabilizes DDX3Y. **(A)** Clone 2 cells were transfected to express FLAG-tagged DDX3X with or without HA-tagged DDX3Y, and cell lysates were processed for IP and western blotting with the indicated antibodies. **(B)** Parental U87MG cells were transfected to expressed HA-tagged DDX3Y, and IP and western blotting were performed for cell lysates. **(C)** Schematic representation of DDX3X and DDX3Y protein seqeunces with lysine/arginine residues of interest marked, generated by BioRender. **(D, E)** Parental U87MG cells were transfected with empty vector (EV) or a plasmid encoding the indicated HA-tagged DDX3Y variant. Representative western blots of cell lysates are shown on the left, and quantification on the right. All experiments were performed in three biological replicates, and results shown are fold change +/- SEM with p values calculated by unpaired student t-tests. **(F)** Parental U87MG cells were transfected to express the indicated HA-tagged DDX3Y variant, followed by treatment with 40 μM MG132 for 6 hours before processed for western blotting. Quantifications of HA band intensity is shown on the right. WT, wild-type, K-R, K250,546R.

Post-translational modifications on lysine residues, such as ubiquitination, are well known to trigger protein degradation (Damgaard, 2021). Among the 32 lysine (K) residues on DDX3X, three (K55, K138, K162) have been linked to ubiquitin-mediated degradation (Wang *et al*, 2021); however, all three are shared with DDX3Y and are therefore unlikely to contribute to the difference in stability between the two proteins. To identify the residues that are responsible for the instability of DDX3Y, we leveraged the high sequence identities between DDX3X and 3Y. Sequence alignment showed two lysine residues on DDX3Y, K250 and K546, which are replaced by arginine residues in the same positions of DDX3X (Figure 4C). We therefore substituted these two lysine residues on DDX3Y with arginine individually. Both the K250R and K546R mutants were expressed at much higher levels than wild-type DDX3Y when U87MG cells were transfected with equal amount of plasmid encoding each protein (Figure 4D), indicating that both residues contribute to the instability of DDX3Y. Combining these two mutations did not further stabilize DDX3Y (Figure 4E). In addition, treatment with proteasome inhibitor MG132 protected wild-type DDX3Y but not the K250,546R mutant (Figure 4F). Together these results suggest that K250 and K546, which are unique to DDX3Y, target this protein to the proteasome-mediated degradation pathway in U87MG cells.

## Discussion

We have observed two different mechanisms through which DDX3X suppresses DDX3Y levels. In HCT116 cells, knockdown of DDX3X results in moderate increases in endogenous DDX3Y mRNA and protein, likely by enhancing *DDX3Y* mRNA stability. However, we cannot rule out addition roles for DDX3X such as repression of *DDX3Y* transcription. In contrast, reduction of DDX3X in U87MG cells does not affect *DDX3Y* transcript levels, but leads to stabilization of DDX3Y protein. DDX3Y is much less stable than DDX3X in U87MG cells, and we were able to identify the two lysine residues that contribute to the lower stability of DDX3Y. Since we also found that DDX3X and 3Y interact with each other, our data favor a model in which DDX3X binds DDX3Y to promote its degradation. DDX3X may do so by recruiting an E3 ubiquitin ligase, but we were unable to detect a decrease in ubiquitination of the stabilized DDX3Y mutants as compared with wild-type protein (not shown). Alternatively, DDX3X may recruit DDX3Y to organelles such as stress granules to accelerate its degradation. Further studies are needed to clarify these possibilities.

Dysregulated DDX3X activity has been linked to a number of diseases, some of which are sexually dimorphic. For example, DDX3X syndrome primarily affects females, whereas Burkitt lymphoma is skewed toward males (Gong *et al*., 2021; Levy *et al*., 2023). In mouse models of DDX3X syndrome, loss of *Ddx3x* in neural progenitor cells leads to increased *Ddx3y* mRNA (Hoye *et al*., 2022; Patmore *et al*., 2020). In light of our results in HCT116 cells, it would be of interest to test if this increase is also due to enhanced *Ddx3y* mRNA stability. However, it is unclear how this increase translates into Ddx3y protein levels *in vivo*, and mouse and human neural development seems to be differently affected by DDX3X mutations (Gadek *et al*., 2023; von Mueffling *et al*., 2024). It is therefore difficult to predict whether a similar compensatory upregulation of *DDX3Y* mRNA also occurs in humans and, if so, has any contribution to the sexual dimorphism of DDX3X syndrome. In B cell lymphomagenesis, DDX3Y protein is detected only in established male lymphoma cells but not in normal male B cells. This ectopic DDX3Y expression has been proposed to contribute to the sexual dimorphism seen in Burkitt lymphoma and other male-skewed cancers with frequent *DDX3X* mutations (Gong *et al*., 2021). Notably, this ectopic expression is probably not caused by an increase in *DDX3Y* mRNA or direct repression of DDX3Y translation by DDX3X, and appears to be a long-term effect that becomes apparent weeks after DDX3X deletion (Gong *et al*., 2021), suggesting that additional alterations are needed. Unlike in HCT116 cells, we were unable to detect the endogenous DDX3Y protein in parental U87MG cells with or without DDX3X knockdown or in Clone 2 (not shown). However, following reduction of DDX3X, a robust upregulation was observed for the exogenous DDX3Y protein expressed from a construct containing just the coding sequence but not the 5’ or 3’ untranslated region (UTR), mainly due to stabilization of DDX3Y protein (Figure 3). Since the 5’ UTR of human *DDX3Y* transcripts can repress translation (Gong *et al*., 2021; Jaroszynski *et al*., 2011), it is possible that additional alterations are needed following loss of DDX3X function to alleviate this translational repression before endogenous DDX3Y protein becomes abundant.

During mammalian sex chromosome evolution, most genes were lost from the Y chromosome but retained in the X chromosome (Bellott *et al*., 2014). Among only 17 human gene pairs that were retained in both X and Y chromosomes, *DDX3X* and *3Y* are unique in that they are extraordinarily dosage-sensitive (Rengarajan *et al*., 2025). In addition, DDX3Y protein is not detectable in a number of male human tissues (Ditton *et al*., 2004), consistent with this protein having deleterious effects such as the oncogenic effect mentioned above. One mechanism that has evolved to keep human DDX3Y protein levels low is the translational repression by AUG triplets in the 5’ UTR in some *DDX3Y* transcripts (Jaroszynski *et al*., 2011). In our attempt to knock out *DDX3X* in U87MG cells, the cells responded to CRISPR/Cas9-generated frameshift by utilizing an originally out-of-frame AUG to restore the open reading frame (Figure 2A). However, DDX3X protein is expressed at greatly reduced levels in this clone (Figure 2B), likely as a consequence of translational repression by the original AUG that is now localized in the 5’ UTR, mimicking the repression of DDX3Y translation that evolved in nature. Two other mechanisms that suppress DDX3Y levels are the destabilization of DDX3Y mRNA and protein by DDX3X, as we describe here. These two mechanisms also protect the cells from dysregulated DDX3X levels by inducing the opposite response of DDX3Y to keep the total DDX3 dosage stable. This protection is apparently disrupted in DDX3X syndrome, as DDX3Y fails to compensate for the loss of DDX3X protein or activity in male patients. Conversely, tumor cells may hijack these mechanisms to ectopically express DDX3Y to overcome the loss of DDX3X. A complete understanding of these mechanisms may lead to new therapeutic approaches by targeting DDX3Y expression.

## Acknowledgements

This work was supported by the US National Institute of Health (R01 DE029802 to S.W.).

## Notes

### Competing Interest Statement

The authors have declared no competing interest.

