## Supplementary data for "Diverse mechanisms of DDX3Y suppression by DDX3X"

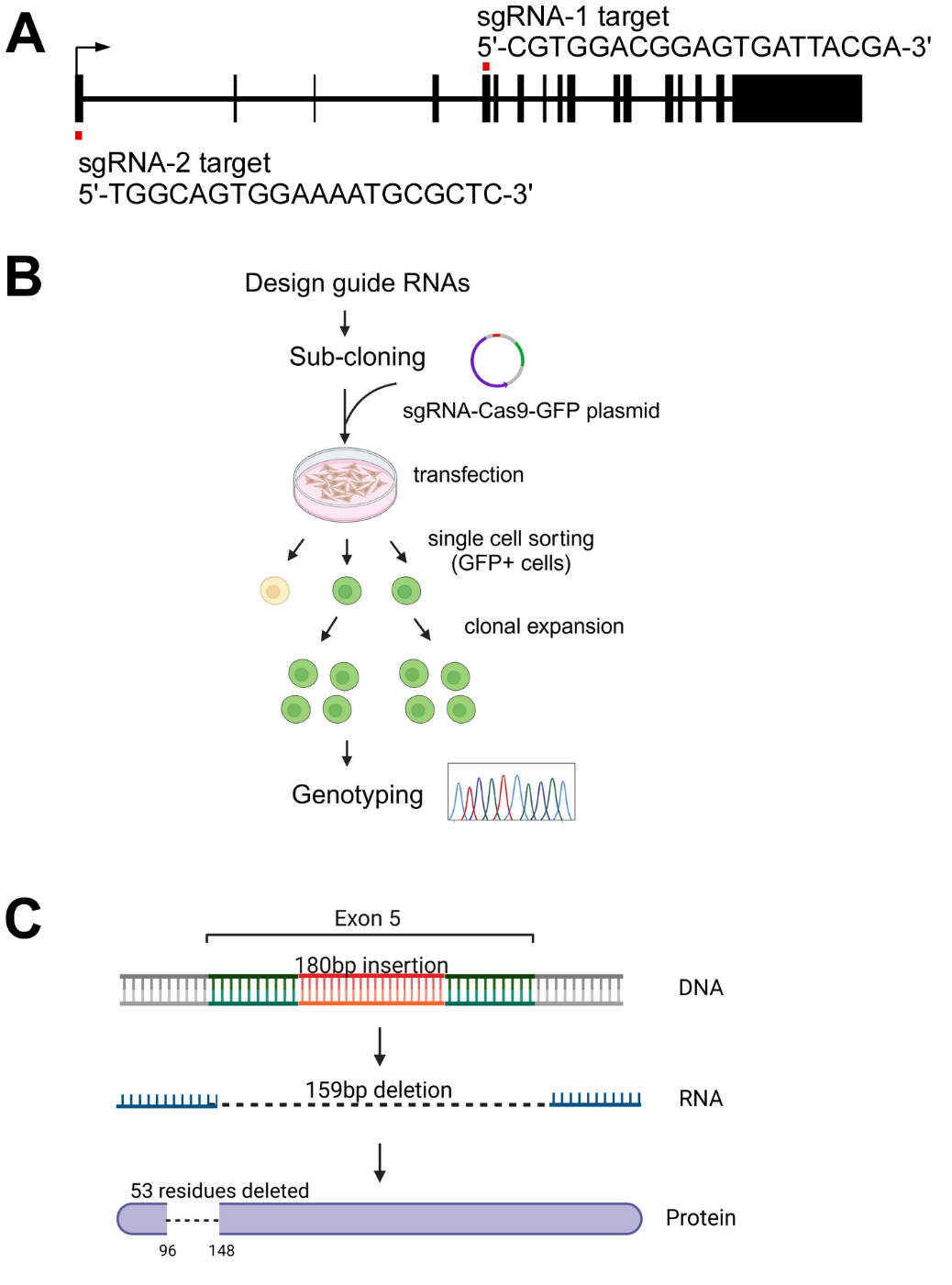

**Supplementary Figure S1. CRISPR/Cas9-mediated gene editing of *DDX3X*.** **(A)** Diagram of sgRNAs targeting *DDX3X*. Black rectangles: exons; red dots: sgRNA target sites; arrow: transcription start site. **(B)** Flowchart of generating U87MG clones with *DDX3X* gene edited by CRISPR/Cas9.

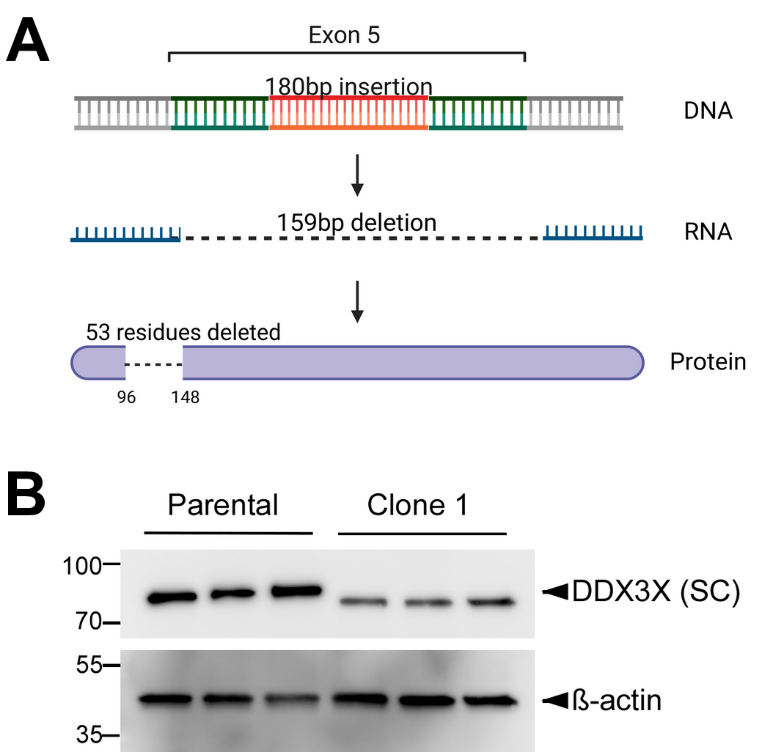

**Supplementary Figure S2. Characterization of Clone 1. (A)** Diagram of the genotyping results of Clone 1 and the putative encoded DDX3X protein. Sequencing of the PCR product amplified from the genomic DNA surrounding the sgRNA-1 target site revealed a large insertion of 180 bp in Exon 5, which resulted in exclusion of the entire Exon 5 (159 nucleotides) as shown by cDNA sequencing, possibly by disrupting an exonic splicing enhancer. As a consequence, the translated protein had 53 residues deleted. **(B)** Western blotting of parental U87MG and Clone 1 cell lysates (three biological replicates each), using the anti-DDX3X antibody from Santa Cruz.

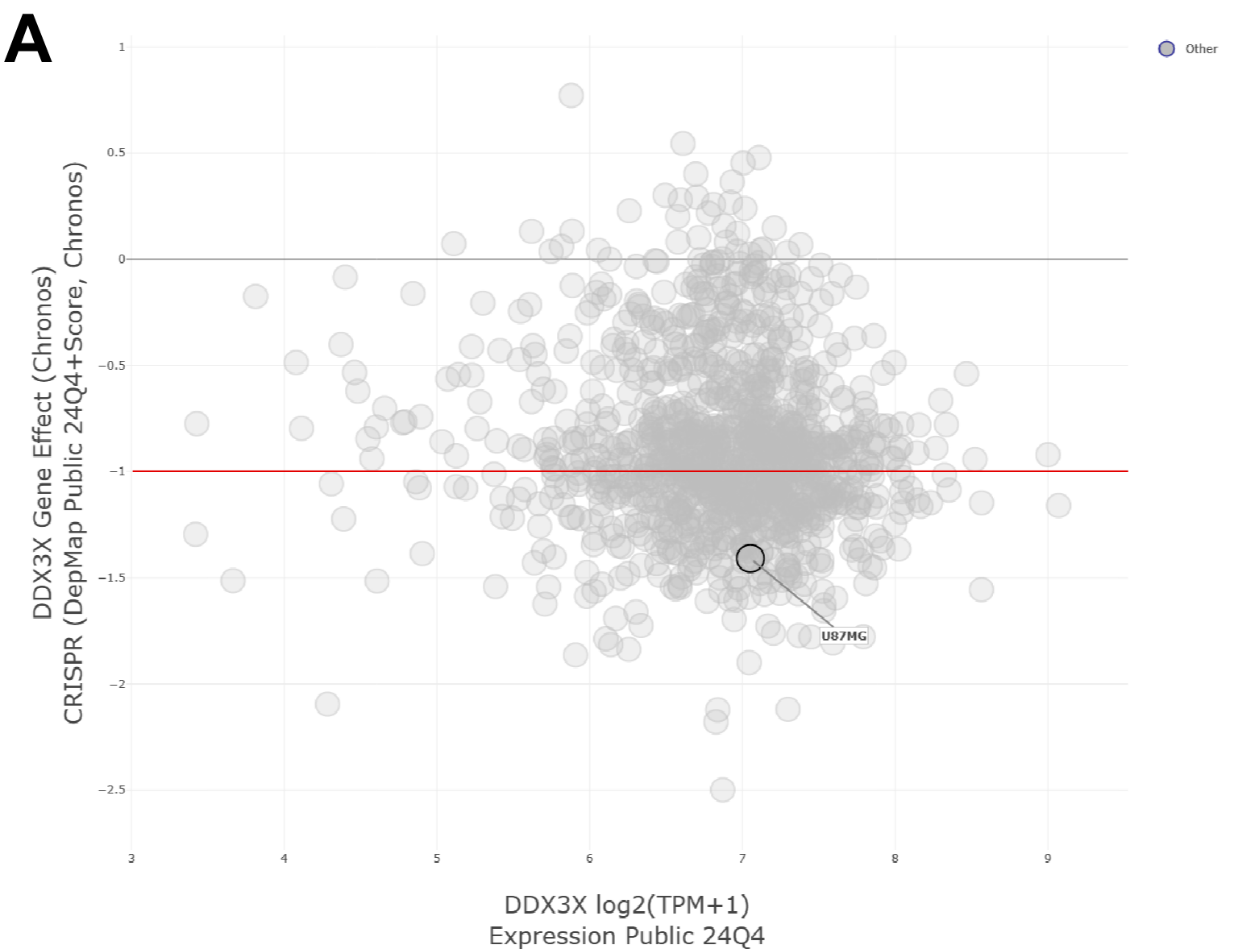

**Supplementary Figure S3. Scatter plot of CRISPR Chronos dependency score vs relative expression level of DDX3X.** Chronos score is representative of gene perturbation effect based on data from a cell depletion assay. A score lower than -1 (below the red line) indicates a high likelihood of a gene being essential for the survival of a given cell. U87MG is circled in black. Plot was generated by depmap.org.

**Supplementary Table S1. Primers (5’ to 3’) used in this study**

**For subcloning into expression vector:**

| *DDX3X*-BamHI-s | GCGCGCGGATCCATGAGTCATGTGGCAG |
| --- | --- |
| *DDX3X*-XhoI-r | AAAAAACTCGAGGTTACCCCACCAGTCA |
| *DDX3Y*-BamHI-s | GCGCGCGGATCCATGAGTCATGTGGTGGTG |
| *DDX3Y*-EcoRI-r | CCGGAATTCAGGTTGCCCCACCAGTC |

**For RT-qPCR:**

| *DDX3X*-s | ACTATGCCTCCAAAGGGTGTCC |
| --- | --- |
| *DDX3X*-r | AGAGCCAACTCTTCCTACAGCC |
| *DDX3Y*-s | CGGCAGTAACTGTCCTCCACAT |
| *DDX3Y*-r | CTGGAGTAGGACGAGTATAGCG |
| *GAPDH*-s | AGGGCTGCTTTTAACTCTGGT |
| *GAPDH*-r | CCCCACTTGATTTTGGAGGGA |

**For CRISPR/Cas9-mediated editing of *DDX3X* gene in U87MG cells:**

| sg1-sense | CACCGCGTGGACGGAGTGATTACGA |
| --- | --- |
| sg1-antisense | AAACTCGTAATCACTCCGTCCACG |
| sg2-sense | CACCGTGGCAGTGGAAAATGCGCTC |
| sg2-antisense | AAACGAGCGCATTTTCCACTGCCA |

**For genotyping Clone 1 and 2 obtained using CRISPR/Cas9 in U87MG cells:**

| sg1-gt-s | TTGTGAGCTGTGTGCCGATT |
| --- | --- |
| sg1-gt-r | TCTGTGTCTTAAAAAGTCAAGCAAA |
| sg2-gt-s | GATCTCGAGAACTCCGAGGC |
| sg2-gt-r | GACAATACAGCGGGCCGAG |

**For cDNA sequencing of Clone 1 and 2:**

| sg1-rt-s | CCAGCAAAGGGCGCTATATTC |
| --- | --- |
| sg1-rt-r | CTTTTGCACTGGAGTTGGGC |
| sg2-rt-s | CTTCGCGGTGGAACAAACAC |
| sg2-rt-r | CAGTTGTTGCCTGTTGCCTC |

**For generating K250R, K546R and double mutants of DDX3Y**

| K250R-s | ATGGTCCAGGAGAAGCTTTGAGGGCTGTGAAGG |
| --- | --- |
| K250R-r | CCTTCACAGCCCTCAAAGCTTCTCCTGGACCAT |
| K546R-s | CCTTGCCACCTCATTCTTTAATGAAAGAAATATGAATATTACAAAGGATTTGT |
| K546R-r | ACAAATCCTTTGTAATATTCATATTTCTTTCATTAAAGAATGAGGTGGCAAGG |
